# IntelliEppi: Intelligent reaction monitoring and holistic data management system for the molecular biology lab

**DOI:** 10.1101/184721

**Authors:** Arthur Neuberger, Zeeshan Ahmed, Thomas Dandekar

## Abstract

Daily alterations of routines and protocols create high, yet so far unmet demands for intelligent reaction monitoring, quality control and data management in molecular biology laboratories. To meet such needs, the “internet of things” is implemented here. We propose an approach which combines direct tracking of lab tubes, reactions and racks with a comprehensive data management system. Reagent tubes in this system are tagged with 2D data matrices or imprinted RFID-chips using a unique identification number. For each tube, individual content and all relevant information based on conducted experimental procedures are stored in an experimental data management system. This information is managed automatically but allow scientists to engage and interfere via user-friendly graphical interface. Tagged tubes are used in connection with a detectable RFID-tagged rack. We show that reaction protocols, HTS storage and complex reactions are easily planned and controlled.

## Introduction

The Internet of things is increasingly becoming a tangible reality of the 21st century with examples from the modern factories for automatic assembly lines that work complementarily to a supportive smart-tag based storage system (1). It is not restricted to certain technological areas, instead, with the addition of some creativity, it can be applied to almost every complex process in which electronic devices are involved (2). It is therefore rather surprising that such a development is still missing in most of the modern biological laboratories, where automation is becoming more and more visible through the innovation, improvement of kits, state-of-the-art omics technology and laboratory information management systems (LIMS).

First attempts to create integrated LIMS can be traced back to the early 80’s in the form of early patents, whereas, real marketing efforts only became noticeable in recent years, when some leading laboratory equipment suppliers started to develop technology-based intelligent laboratory systems. A good example for this development is the advent of fully automatic systems like Eppendorf’s robotic workstation “epMotion” (3) or Dornier-LTF GmbH’s pipetting robot “PIRO” (4). An automated process for analysis has been proposed in several patents. For instance, Lang and colleagues (5), included “storage” of experimental lab data in the memory of a contactless chip card or a barcode attached to the corresponding reaction vessel. Moreover, RFID (Radio-Frequency Identification) technology has already successfully been implemented commercially in the electronic identification and administration of blood donations in transfusion medicine (6, 7).

Even though lab machinery and devices work quite efficiently within their own technological boundaries but almost no integrative interconnection between these smart tools has been established to date. These “closed” systems are unfortunately not yet smart and integrated enough to fully replace the operator and his manual interventions in an environment like the academic molecular biological lab. This demand for facilitated and smart, yet partly manually controlled half-automation of experimental processes could only be met in a semi-open system. Most of the available sample data management systems lack full integrity, i.e. a smart solution that would integrate common devices used on a daily basis, like pipettes and test tubes, into a semi-automatic system.

Facing this stagnation of integrative innovation in the laboratory equipment market, we propose a holistic system that we call “IntelliEppi” (8). It brings the concept of internet of things to the labs. The system is a proof of concept for an internet of things lab management, which, combines economic and easy tagging via two-dimensional data matrix codes or printable RFID tags, with a smart reagent tube logistics and experimental data management software. It supports tracking of all components can be tracked with the help of IntelliEppi, enhance to deliver reagents on time, at the right place, allowing complex synthesis processes and improved quality in the process and its controls.

IntelliEppi allows to store and modify molecules by monitoring, guiding and storing reaction vessels in defined positions, using a reader-system (via a matrix printer or RFIDs) and the power of a versatile program code. It leads to better quality, reproducibility, and transparency in biochemical and molecular biology experiments (9). It incorporates product data management (PDM) from the start and allows reaction tube lifecycle management including long term storage processes, tracking of resources and tubes, and a scheduler. Using IntelliEppi simple reactions and really complex omics experiments can be planned and different tagging strategies in the lab can broadly be explored. All this is made available with the hope to promote the vision of an internet of things from the test tube to the laboratory scale, bundled here in the concept of the intelligent reaction vessel monitoring and actually doing all reaction steps in various experiments. We tested all ingredients and make them available, which includes software, different tagging strategies, performance data, tutorials, labels as well as pseudocode and examples on the synthesis processes.

## Results

### IntelliEppi components and usage

The system IntelliEppi is holistic and comprehensive to provide an internet of things for the molecular biology laboratory and the whole life cycle of the reagent tube:

1. Reagent tubes in this system are tagged with cheap, yet resistant printable plastic RFID chips or 2D data matrices that are marked (e.g. using a laser) onto the top of their lid. These tags assign and store a unique identification number through which each individual tube’s content and all relevant information on conducted experimental procedures of the sample inside the tube can be requested on demand and instantly gathered (Fig. 1(A-C)).
2. All information is stored in a database that is managed semi-manually by the scientist. A user-friendly GUI allows constant information gathering and data editing.
3. Tagged tubes are stored in a smart tube rack which can be tracked via its own RFID - tag. This happens both physically via long-distance scanning and digitally with all relevant information connected to the tag displayed when a specific sample tube needs to be found and identified.
4. The rack can be scanned for information on tube contents and the place of the tube inside the rack. Alpha-numerical - or colour-coding on the rack assign each tube an individual slot on the rack.
5. The rack itself, inside the fridge for instance, can be easily identified via a lighting up LED, which is activated when it is scanned. This way, no samples are lost due to unknown location or erased labels. This makes tracking of old samples and connected information easier and faster.
6. The database stores all relevant information for individual tubes, entire racks, and chemicals (buffers, reagents etc.). All components are stored in tagged tubes or flasks and can therefore be integrated via their identifiers into an interactive digital experimental protocol. This allows both precise design of an experiment and guidance through it, with possible alterations by the user when conducting the experiment. Here, performed steps and other relevant information are updated to the database and stored for every individual tube.

**Figure 1.**
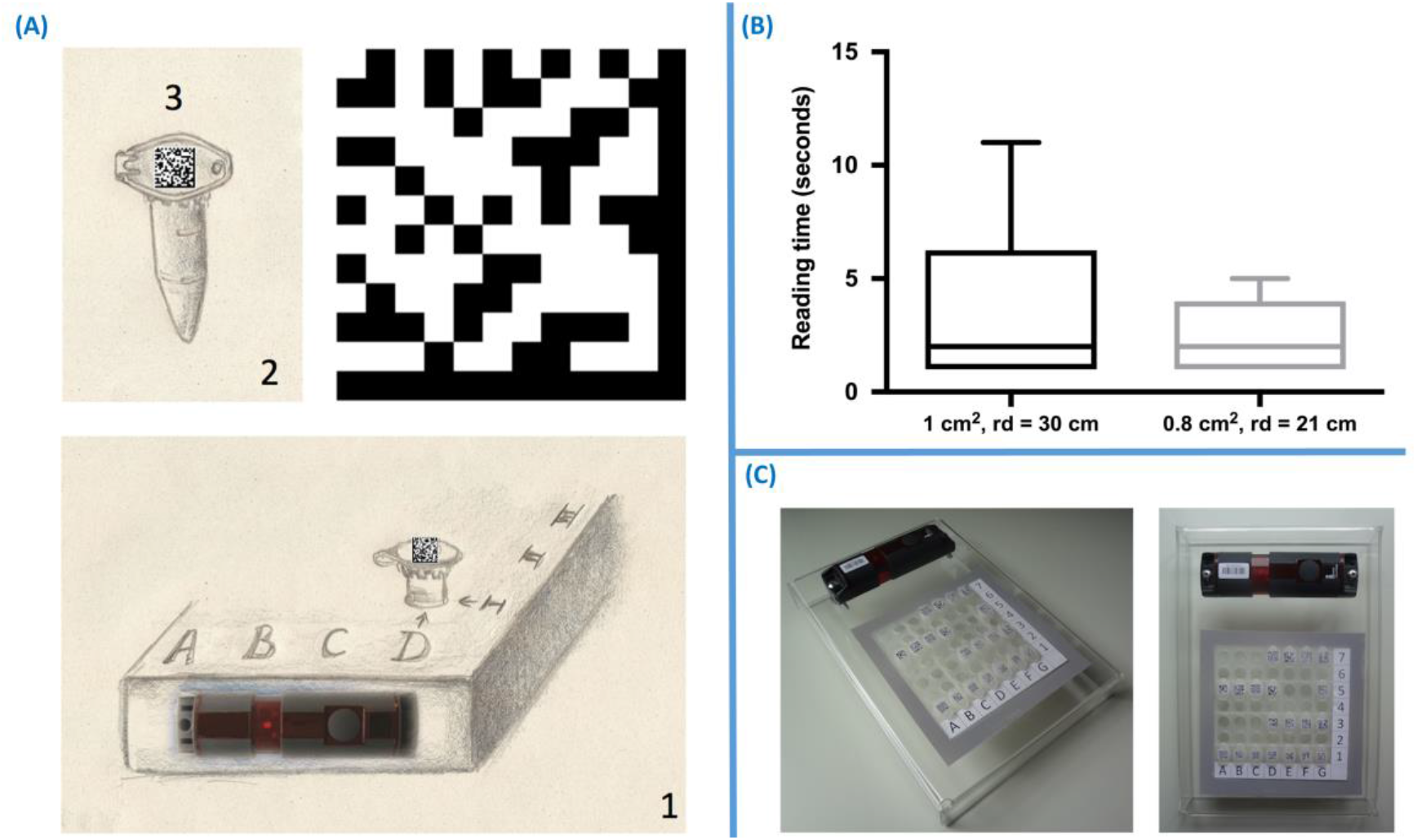
**(A)** Tagged SMARTtube, (B) box plot of scanning time data matircs, and (C) SMARTrack prototype.

### IntelliEppi workflow and validation

IntelliEppi’s power is demonstrated with the aid of conceptual coding examples for different chemical reactions. The combination of software, matrix code or RFID guiding and intelligent tubes achieves an equivalent of NCR guided technology for biochemical reactions.

A standardised process flow is schematically outlined in Fig. 2(A), individual steps are shown in Fig. 2(B). As an appropriate practical example for a complex synthesis we have chosen the activity and kinetics real-time monitoring of the T4 polynucleotide kinase reaction. It is based on a singly labelled DNA-hairpin smart probe coupled with λ exonuclease cleavage (12) (our demonstrator model system). In their approach, Song and Zhao designed a smart probe (labelled oligonucleotides with a hairpin shape) with a fluorophore at the 3’-phosphate end. The Fluorescence is quenched by a guanine-triplet at the terminal 5’-hydroxyl group (12). In the presence of ATP, this 5’-hydroxyl group of the smart probe is then phosphorylated by the T4 polynucleotide kinase (Fig. 2(C)). In a second step, fluorescence enhancement is caused when the resulting 5’-phosphoryl end is cleaved by the λ exonuclease. The latter is a 5 → 3 exonuclease with a preference for phosphate moieties at the 5’ of double-stranded DNA ends that degrades these double-stranded DNA while yielding mono-nucleotides and singlestranded DNA. The detection process is carried out in a real-time PCR instrument as this offers smooth temperature control, sealed tubes, and high-throughput detection (12).

This experiment can be divided into various steps (Fig. 2(B)), each of which is associated with important information. In the first stage, a test tube is tagged with an individualized 2D barcode using IntelliEppi’s “Generator” function. When using the latter, the user defines the experiment, e.g. “T4 polynucleotide kinase (pnk) reaction", and manually fills out the desired range of accompanying options, such as the conducting researcher’s name, the SmartTube’s No. as well as the SmartRack’s No. and SMARTtube storage position within this, and the date and time of the experiment. By clicking on the 2D barcode symbol, the user generates a 2D data matrix which stores the information from all fields in a compact, yet comprehensible code format like “ST73|SR02|A5|T4 pnk reaction|19/07/2016 19:30:15". The latter, being encoded in the actual 2D tag label, can later be easily read for identification of the SMARTtube, e.g. in a rack inside a fridge. A further click on the database symbol stores the entire information package into the ExKPDM database for long term storage, identification and data management in the future.

Fig. 1(A) illustrates the SMART reaction control. Template DNA and a nucleotide mix are then added to the tagged tube (Position 1). If preferred by the user, this step and the following ones are manually noted down in the corresponding ExKPDM file that was created for the registered tube: In position 2, the T4 polynocleotide kinase is added and therewith the reaction started. The reaction is stopped after 30 minutes (Position 3). The now labelled reaction can then be added to the hybridization platform and maintained there for a total reaction time of 24 hours (Position 4) before the filter is taken out (Position 5) and transferred to the detector for read out (Position 6).

IntelliEppi is able to perform the described experiment whilst producing an up-to-date and lasting live documentation. A single document file is semi-automatically updated at every step in the reaction flow. After the experiment, the tube (containing the product of the T4 pnk reaction) can be tracked via its 2D tag using a scanner connected to the ExKPDM system (run on a computer; scanner connected via USB boost or Bluetooth; more details on tagging and tracking suppl. Material, part 1). The T4 polynucleotide kinase reaction SMARTtube can be stored on a SMARTrack which is also registered in the system under the same experiment. However, in this case, the SMARTrack’s RFID tag is scanned by an RFID reader. The latter is also connected to the ExKPDM system. The reader is, as it is the case for the optical 2D data matrix scanner, connected via USB boost or Bluetooth with a computer. Identec Solutions’ system works with TCP/IP or COM connections. A reading device is currently under development that can be directly connected to a computer (via USB boost).

ATP, the smart probe, T4 polynucleotide kinase, λ exonuclease, and other reagents used in various experiments in this lab can be stored together in designated SMARTrack. The following features are conceptual and under current implementation into the software package: Reagents, buffers, and other chemicals could also be registered in the ExKPDM chemical database. All chemicals (reagents, buffers etc.) would have their own individual ID/EPC. Hence, when a new experiment is designed, using the experimental protocol function of the ExKPDM, all experimental components (buffers, reagents etc.) as well as the SMARTtubes, which these experiments are about to be performed in, would be manually identified using a search query and then registered into the new experimental protocol. In this protocol, all experimental steps, e.g. how much ATP shall be put in at which time point into which SMARTtube, is clearly pre-defined before any action is taken. This protocol could also be manually adjusted at every point of the experiment (as long as the user has the rights to do this). In case of experiments in which certain timing points must be observed, a timing function enables the user to plan this dimension of his experiments as well.

**Figure 2.**
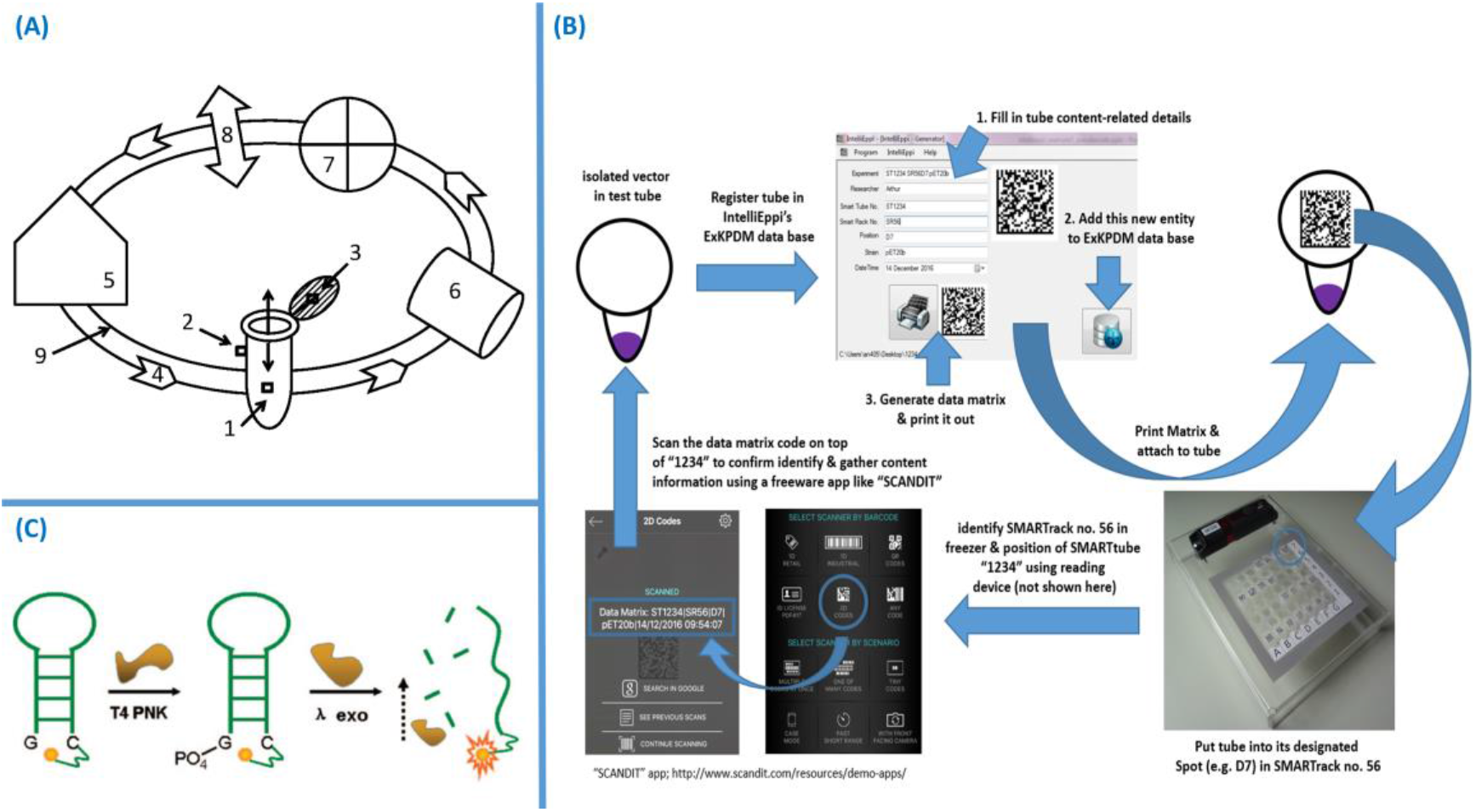
(A) The reaction cycle, (B) summary for one example of a simple application routine for the IntelliEppi system, and (C) T4 Reaction at the terminal 5’-hydroxyl group.

As previously mentioned, such a digital protocol would be interactive, changeable at any point and more like a programme that is running in parallel until it reaches its defined end. This programme would automatically notify the user when it is time for him to execute a timing-dependent step in the experiment. It would also provide information on the next step to execute. In the era of smartphones and tablet PCs, one might even consider a complementary app that informs the user everywhere and at any time. This way, experiments can be easily managed from abroad, or outside the lab, by an instructor or group leader. Every change of a SMARTtube’s content is written into the experimental protocol and subsequently into the SMARTtube database. Next time when this SMARTtube is scanned, the user receives the updated information as well as the full experimental history of this tube (i.e. all executed steps, content changes together with the corresponding time points and maybe also the name of the person who conducted this step).

The internet of things for the laboratory requires integration of software and lab components. Thus, besides tagging and tracking for programming of reaction courses and integration of all components, the software is critical. SMARTtubes belonging to the same experiment or analysis fraction are always stored in the same corresponding SMARTrack, except during their handling, i.e. when the content of a tube is changed or analysed for instance. The IntelliEppi laboratory database, as one of several software components of the ExKPDM system, stores the EPCs/IDs of each SMARTtube and - rack. The user who is trying to find an individual tube for further experimental procedures could either directly search for the tube by typing the ID key or EPC into a search tool, or, alternatively, search for the corresponding SMARTrack (in which this tube was stored) using the SMARTrack-ID. As an output of such enquiry for an individual SMARTtube, or a whole SMARTrack, an interactive window opens that displays all information concerning this SMARTtube or, in the case of a search for a whole SMARTrack, all registered SMARTtubes belonging to this rack, respectively. In the latter case, manual picking of an individual SMARTtube (via SMARTrack window) would display the lab history of this tube (as it is also the case for the output of a manual search for this SMARTtube).

Fig. 1(C) illustrates a hypothetical mobile version of IntelliEppi. Information was analysed on the position of the searched tube in this SMARTrack (row and column, e.g. “D1”); the place where (in which fridge, floor etc.) the SMARTrack was located (e.g. “Fridge II, Lab. 3, 2nd Floor”); the time when the sample had been created (e.g. “Experiment started: 17:10:00 04.07.2013”). The system considers also the experiment this SMARTtube belongs to (e.g. “T4 pnk activity”), when and how the content was changed. All this is analysed via an interactive experimental history sliding window (see “Add 0.5 mL Buffer 17:18:23 04.07.2013”) using a user-friendly GUI. Cross-links lead to additional information: e.g., on the experimental protocol (created by the user using protocol function of ExKPDM software), on chemical knowledge of the reagents used for analysis (see stylised selector: “Go to chemical knowledge base”) or content change/protocol following, on EPCs/IDs and technical information of lab equipment used in connection with this tube or its content respectively.

A tube´s assigned location in a rack can be easily and economically controlled using colour - and/or numerically coded rack surfaces. For example, an individual tube could be allocated a pre-defined storage position like D1 or red/D or red/1 in the SMARTrack. Columns and rows are thus labelled and/or colour-coded in the rack. Obviously, a user looking at a SMARTrack filled with SMARTtubes (datamatrix coded for example) cannot possibly distinguish between the tubes on the basis of different codes on the top of lids. Thus, it is required that every individual tube, after being taken out of the rack, e.g. for centrifugation or pipetting, is always put back in its pre-defined position in the rack, like D1. Information on which the right place for an individual tube is, could be instantly recalled by scanning its tag.

### Designing complex reactions with IntelliEppi

More complex reactions often involve molecular biology genetic engineering. As an example (Fig. 3 (A)), a fusion construct consists of PCR generated fragment with a light-gated BLUF domain is attached to a T4 polynucleotide kinase. This makes the T4 polynucleotide kinase light-dependent and so light can be used as an additional external control in the system. The final construct is stored in position end.

**Figure 3.**
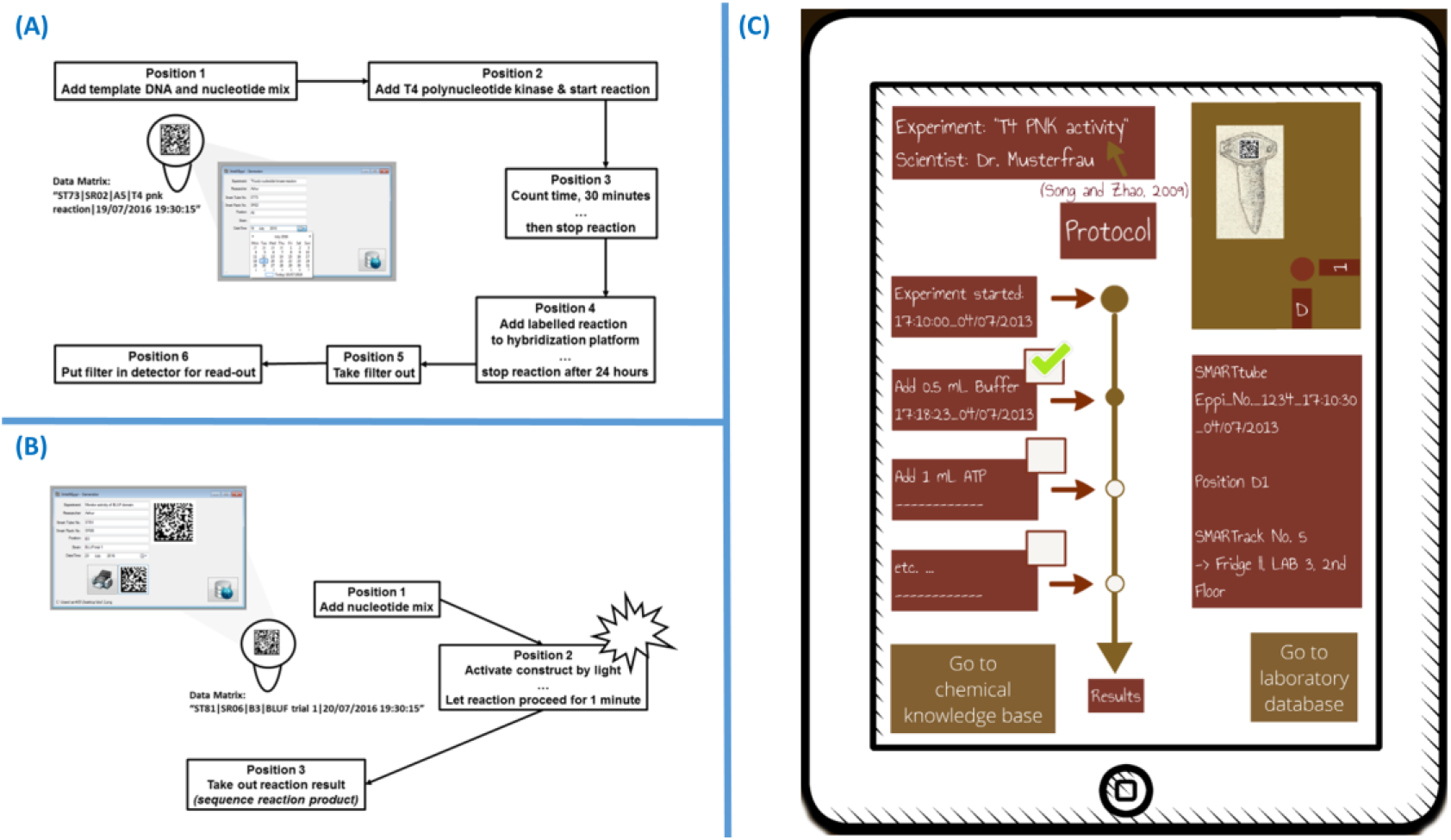
(A) The T4 polynucleotide kinase reaction is given as an example for chemical reaction pathway engineering using IntelliEppi. (B) BLUF domain fusion activity testing as an example for chemical reaction path engineering using IntelliEppi. (D) Abstract mobile IntelliEppi-ExKPDM user interface.

This more complex program consists of the following subroutines: Cloning of BLUF domain (Fig. 3(B)), 1: Template DNA extraction from *E. coli*, 2: Add suitable PCR primers; PCR the BLUF domain, 3. Easy cloning step to get BLUF domain in vector of choice. 4: Template T4 DNA, PCR the T4 polynucleotide kinase, 5: Easy cloning step to get T4 kinase domain in vector of choice, 6: Transform the vector into *E. coli*), and Expression and purification of the BLUF domain (Fig. 3(B)), 7: Express the fusion construct, and 8: Purify the fusion construct on a nickel column). Finally, the activity of the new construct is tested in an example where the IntelliEppi helps to monitor the underlying chemical reaction pathway.

Another area where the additional smart control and the IntelliEppi system with software is handy is for complex synthesis processes. Examples explored here are dendrimer synthesis, a DNA macrame or RNA designer aptamers wired for a logical gate or array.

## Discussion

Recent alternatives to IntelliEppi’s software and monitoring components include the pipetting robot Pirot as a bench worker robot and the Emerald cloud where you order a service with a programming language that helps to manage different types of laboratory work. Several further software languages allow to formulate experiments. For instance, there is Biocoder, a programming language for standardizing and automating biology protocols (11). The PaR-PaR laboratory automation platform (12) features the “biology - friendly high-level robot programming language PaR-PaR (programming a Robot; http://prpr.jbei.org/). Furthermore, there is the formalization language EXACT (13). Moreover, you can order experiments from Emerald cloud even though there is no automation for this. However, these are only partial solutions. We provide an integrated solution: IntelliEppi complements the efforts described above by allowing the detailed design and step-by-step monitoring of complex biochemical laboratory experiments and by using a specific programming language focused on position and function codons for guiding and monitoring the reaction path together with the reaction vessel (the “Eppendorf tube”) in an intelligent way. A less technologically advanced but also much cheaper and “semi-open” approach for half-automated sample management is nowadays realized in the use of label printers for individual samples (14). In some cases, these printers are sold with basic sample data management software (e.g. Brady Laboratory Labels).

The identification of molecular biology laboratory samples in a container using RFID - technology, has been proposed by Excoffier and colleagues (15). A combination of barcode and RFID tagging on the same test tube has also been proposed in form of a system where test tube identification data is encrypted in a barcode whereby additional data is stored on the RFID tag (16). An advancement of the latter is a system comprised of a rack holding element and, adjacent to this, of a carrier for an antenna for the wireless reading of an RFID tag attached to a test tube (17). In a different, further automated system, test tubes, tagged via barcode, are transported in a RFID-tagged conveying device via a conveyor belt to pretesting, testing, and post-testing stations. Corresponding devices are mounted to the system whereby identification and control of action in this system are based on RFID tagging of the conveying devices (18). With a view to relevant patents for RFID technologies, printable RFID transponders are likely to constitute one of the most promising approaches (10). The corresponding printing technology has already been proposed in a separate patent (19). This is only an excerpt from the lively patent scene for RFID - and barcode-based life science applications.

Some proposed approaches aim to combine tagging with simultaneous content analysis or reaction monitoring, like those using encapsulated microprobes and lab-on-chip systems. The latter particularly apply to diagnostics. A system-on-chip digital pH meter, for instance, has already been successfully enclosed in a diagnostic capsule for in vivo diagnostics of gastro-intestinal conditions and diseases (20). An even broader spectrum of analytical measurements is offered by a micro-lab system that is encapsulated in a biocompatible shell (21). Digital cloud-based lab management systems (Laband.me LTD is a good example for such a provider; see also https://www.laband.me/) offer easy-to-use data recording and analyzing tools, based on apps, cloud-storage, and electronic notebooks, making the daily management, simple statistical analysis, or straightforward visualization of data easier. However, none of the mentioned systems integrate system tagging, detection and analysis into an intelligent and user-friendly management software solution.

Our software has the advantage to be easily, generically adapted to new, different biochemical experiments and is easy to be scaled up. Given examples include the T4 polynucleotide kinase reaction cycle, a more complex genetic engineering example with PCR protocol, and some complex synthesis examples. Furthermore, the tool also enables a library generation for RNAseq, or omics-seq experiments, ClipSeq, ChIPseq, RIPseq. These protocols are not shown here but are easily accomplished through this strategy, which affords relative flexibility to the reaction tube position, typical reaction steps as well as storage and different addition modification and filtering steps.

IntelliEppi has to be seen as a complementary part of a general effort to automate, monitor, and design lab experiments. This includes the Emerald Labs where you can order experiments from these. There is also the use of a robot scientist, well-known from an early article (22), (23). The aim is again to free the mind for intelligent work. There is also the idea to increase the flexibility in experimental design (as evidenced by our flexible code) pursued by current efforts, for instance in instructing 3D printers (24), flexible synthesis-like software (25) and HTS advances such as digital picoliter PCR(26), (27).

This is a first step in the direction towards a micro-factory environment. Further extensions include the synthesis of even more complex compounds and further miniaturization.

## Methods

### SmartTube and SMARTrack

Test tubes (e.g. provided by the Eppendorf) are involved in highly complex processes conducted in a modern biochemical or molecular biological lab. It is therefore evident that these tubes act as the cardinal element in a lab logistics system. Most laboratories are still required to manage their sample and data logistics by hand. Thus far, the most innovative sample data management system to mention is based on label printers that are capable of creating labels of alphanumerical content as well as 2D codes which have a sticky back via which these can be attached to tubes. In a given situation, as shown in Fig. 2(A); a tagged reagent tube’s (“SmartTube”) lid surface was tagged with a data matrix. SMARTtube (sitting inside a SMARTrack) tagged with a data matrix ECC encoding “ASCII” (ECC 200 for instance). SMARTrack, tagged via i-Q350L FLSensorSMART, with a SMARTtube inserted at a designated position. SMARTtube tagged with ECC200-datamatrix. ECC-datamatrix encoding “ASCII”.

### Datamatrix coding

An alphanumerical code like “ST1234SR56D7 17:10:38 03.09.13” for instance can be encoded in a data matrix composed of 20 x 20 modules. This code can be interpreted as: “SMARTtube no. 1234 in SMARTrack no. 56, positioned in row D and column 7 in this rack, last procedure was executed at 17:10:38 on 03.09.2013". A module size of 0.4 mm leads to a total code area size of 8.0 mm2. This dimension is small enough to be lasered onto the top of a standard reagent tube lid. We tested the data matrix coding. It is large enough to be read from the maximal distance of 30 cm (for a 1 cm2 matrix; 21 cm in case of a 0.8 cm2 matrix) within 2 seconds on average using a simple freeware smartphone app (Fig. 2(B)).

### RFID-tagging

In the case of a RFID tag printed on top of a tube lid, its electronic product code stored in the memory of this RFID tag (that enables individualisation of every single tube) could be read by an appropriate RFID reader device (depending on the operating frequency in Hz). For this technology, we tested RFID tags made by Identec Solutions AG. These are moderately sized RFIDs (6-10 cm) that are commercially used for location tracking of lorries and are utilized in our system for reaction tube rack tracking in a laboratory setting. In combination with a read-out device, these RFID tags were obtained from Identec Solutions AG. Anticipating future developments, we also looked at the RFID potential from PolyIC’s imprintable RFID tags. Here, different polymer foils are coated with electronics - potentially a very powerful technique that would enable submillimetre miniaturization of cheap tags for reagent tubes. However, this technology is not yet in production mode. According to the developing company, this technology will take another 3-5 years until it becomes available. Hence, the best technology currently available features 3-4 mm RFIDs from Microsensys. As mentioned earlier, 2D coding was used in our demonstrator model. Here, an individual tube is allocated an individual ID key that is stored in an IntelliEppi database as a part of a whole Experimental Knowledge and Product Data Management Software.

### Demonstrator Model

For our demonstrator model, data matrices were generated using the 2D barcode “Generator” module of the IntelliEppi software. These barcodes have the following meta data: Experiment, Researcher, Smart Tube Number, Smart Rack Number, Position, Strain and Data Time. SMARTtubes are always used in a logical connection with an RFID-tagged rack, henceforth referred to as “SMARTrack”. Here, Identec Solutions GmbH, for instance, provided a battery-powered active tag, i-Q350L FLSensorSMART (Fig. 2(C)) for our model system. This tag operates at 868 MHz (EU-compatible) with a localisation and a read/write range response mode of several hundred meters in free air, a memory size of 10,000 bytes (user definable), a 48-bit fixed identification code, a replaceable Lithium battery, and a special marker function with an operating frequency of 125 kHZ. Tag dimensions are 137 x 37.5 x 26.5 mm and therefore suitable for placement at the short side of common tube racks. Furthermore, an LED responds to tag scanning with a bright optical signal when the tag is detected in our model. One can imagine that dozens of SMARTracks are stored in a fridge at the same time. Scanning the content of this fridge for SMARTrack no. 56 for example, gives back an optical signal that allows quick identification of the right rack (where the searched tube is stored). SMARTtubes and the SMARTrack, in which these SMARTtubes are permanently stored, form an intelligent unit within the IntelliEppi system. Together with a reader (and scanner) they form the hardware component of the IntelliEppi system.

### Experimental Knowledge and Product Data Management Software

This is henceforth referred to as “ExKPDM”, is the software component of this system. The ExKPDM software components are shown in Fig. 3(A and B) as a conceptualized work-flow overview of basic components, their cross-linking and interaction within IntelliEppi, and the resulting user benefit. Labels are as follows: 1: RFID tag as a reusable probe insight the tube; 2: RFID tag attached to or a 2D data matrix barcode printed/lasered on the surface of a tube; 3: RFID tag incorporated into or a 2D data matrix barcode printed/lasered on the lid of a tube (incorporation as part of manufacturing). 4: the process flow (not necessarily cyclic); 5: storage module; 6: reaction module; 7: detection module; 8: “mini-factory”, semi-open system (user intervention is possible) for complex syntheses; 9: ExKPDM including IntelliEppi - Tracking. (B) ATP + 5’ - dephospho - DNA -→ ADP + 5’ - phospho - DNA (Figure adapted from (12)).

Specific software modules are developed according to user needs. The user feeds the ExKPDM system with instructions or data via a user-friendly user interface. All further processes run based on this interface in the background, not visible for the user as such, Fig. 3(C). Briefly, individual SMARTtubes are operated through the O-module. Data on each SMARTtube is saved into the laboratory database D via the module for optimal SMARTtube management. The chemical knowledge base enables the user, via the user interface to carry out a system-integrated search in various chemical databanks. The data are also transferred into the laboratory database ingtegrating query results and programmed searches. Final results of the search module appear via the user interface. Finite working cycle (life cycle) data can be requested using 2D data matrix/RFID tagging of SMARTtubes and SMARTracks linked to the ExKPDM.

### Code availability

The executable of the ExKPDM system and a test database are available. The source code will be made available in the same way upon acceptance of the manuscript without any restrictions of usage. Tagging, protocols, RFID information, software and tutorials are all available at: https://www.bioinfo.biozentrum.uniwuerzburg.de/computing/intellieppi Furthermore, source code, setup and executable are all loaded up to the GitHub public repository at https://github.com/drzeeshanahmed/IntelliEppi

## Author contributions

A.N. did all work on the IntelliEppi Lab implementation including demonstrator, RFID tests, data matrix tests and tagging. Z.A did all work on the software aspects of IntelliEppi including the expert system and implementation of code. T.D. lead and guided the study, analyzed the performance and data of the IntelliEppi system. All authors drafted the manuscript and finalized it together.

## Acknowledgements

We thank BMBF for support (grant number 031L0129B).

